# Differential formation of motor cortical dynamics during movement preparation according to the predictability of go timing

**DOI:** 10.1101/2023.07.20.548512

**Authors:** Soyoung Chae, Jeong-Woo Sohn, Sung-Phil Kim

**Affiliations:** Ulsan National Institute of Science and Technology, 50, UNIST-gil, Eonyang-eup, Ulju-gun, Ulsan, 44929, South Korea; Catholic Kwandong University, 24, Beomil-ro 579 beon-gil, Gangneung-si, Gangwon-do, 25601, South Korea

## Abstract

Motor cortex not only executes but also prepares movement, as motor cortical neurons exhibit preparatory activity that predicts upcoming movements. In movement preparation, animals adopt different strategies in response to uncertainties existing in nature such as the unknown timing of when a predator will attack—an environmental cue informing ‘go’. However, how motor cortical neurons cope with such uncertainties is less understood. In this study, we aim to investigate whether and how preparatory activity is altered depending on the predictability of ‘go’ timing. We analyze firing activities of anterior lateral motor cortex (ALM) in mice during two auditory delayed-response tasks each with predictable or unpredictable go timing. When go timing is unpredictable, preparatory activities immediately reach and stay in a neural state capable of producing movement anytime to a sudden go cue. When go timing is predictable, preparation activity reaches the movement-producible state more gradually, to secure more accurate decisions. Surprisingly, this preparation process entails a longer reaction time (RT). We find that as preparatory activity increase in accuracy, it takes longer for a neural state to transition from the end of preparation to the start of movement. Our results suggest that motor cortex fine-tunes preparatory activity for more accurate movement using the predictability of go timing.

**Significant statements:** Anticipating when to move is important in movement preparation. However, it is unclear how motor cortex prepares movement depending on how easy that anticipation is. To answer this, we examine motor cortical activity of mice during a delayed-response task. While motor cortical activity rapidly reaches a “movement-ready” state with unpredictable timing of a go signal (go timing), it does so more gradually when go timing is predictable. Moreover, when go timing is more predictable, motor cortex produces more accurate movement with, unexpectedly, a longer response time. This suggests that rodent motor cortical neurons can resource time information.

## Introduction

Movement is prepared before its execution, and is often initiated by environmental cues instructing ‘go.’ The firing activities of motor cortical neurons during movement preparation are referred to as “preparatory activity” (Vyas et al., 2020). In the dynamical system view, the final state of preparatory activity right before the go cue (i.e., the preparatory end-state) determines the subsequent neural dynamics of movement execution. As such, ensuing behavior (e.g., reaction time (RT) and arm speed) can be predicted from the preparatory oend-state (Churchland et al., 2010; Afshar et al., 2011). In this regard, preparatory activity has been understood as a neural process to achieve an appropriate preparatory end-state to produce desired movement (Churchland et al., 2006b; Churchland et al., 2010; Churchland et al., 2012; Shenoy et al., 2013; Michaels et al., 2016).

In movement preparation, it is important to predict when to move. For instance, responses are faster and decisions are more accurate when a go cue is given at the expected timing (Jaramillo and Zador, 2011). However, little is known about how motor cortex leverages this temporal prediction to enhance behavioral performance and whether the availability of temporal prediction influences movement preparation strategies.

Previous studies have shown that when go timing is predictable, the time left until go timing influences how quickly preparatory activity progresses towards the end-state (Kilavik et al., 2014; Murakami et al., 2014). On the other hand, when go timing is unpredictable, preparatory activity quickly reaches the end-state and sustains it until the go cue is given (Inagaki et al., 2019). However, whether and how the predictability of go timing affects the dynamic profile of the preparatory activity is unknown.

Therefore, we aimed to study whether the dynamics of preparatory activity in motor cortex alters according to the predictability of go timing and if so, how the altered dynamics copes with both unpredictable and predictable go timing. We analyzed firing activity of ALM while mice performed one of the two different auditory delayed-response tasks: random-, or fixed-delay task. In the random-delay task, delay length was randomly given across trials for unpredictable go timing, whereas in the fixed-delay task, prediction of go timing was enabled via fixed delay.

If the preparatory end-state – which reflects the dynamics of preparatory activity – is identical between the two tasks, ensuing behavior should be similar. However, we observed differences in behavioral accuracy and reaction time (RT) between the tasks, as well as in the degree of selectivity at the end-states. These results indicate that the dynamics of preparatory activity vary based on the predictability of go timing. We selected and examined a putative component of ALM population activity assumed to represent movement preparation specifically. In the random-delay task, the preparation-specific component largely featured “readiness to move,” exhibiting negative correlations with RT. In contrast, in the fixed-delay task, preparatory activity reached the end-state at a later stage of the delay. The preparation-specific component was found to have positive correlations with behavioral accuracy, demonstrating motor decision. Intriguingly, the component also showed positive correlations with RT, exhibiting a similar pattern to the positive correlation between RT and accuracy (speed-accuracy trade-off). Our results suggest that motor cortex can fine-tune its firing dynamics during movement preparation based on given temporal resources.

## Materials and Methods

### Behavioral tasks

In this study, we analyzed the open dataset of 11 mice (aged from postnatal day (**P**) > 60). The datasets were made publicly available by the Svoboda laboratory at FigShare (https://doi.org/10.25378/janelia.7489253). A detailed description of data collection procedure can be found in Inagaki et al. (Inagaki et al., 2018; Inagaki et al., 2019). In brief, the mice were trained to learn the auditory delayed-response task (Fig. 1). Five mice were trained to perform the auditory delayed-response task with randomized delay lengths – the random-delay task. At the beginning of the trial, an auditory stimulus was presented at one of two frequencies, 3 or 12 kHz, which informed the water port location to the right or left, respectively. After 1.15 s of presenting the auditory stimulus, a delay of a certain length was given. Then, given a non-selective auditory go cue, the mice had opportunity to lick right or left based on the given auditory stimulus. Delay lengths were randomly selected from eight values: 0.3, 0.5, 0.7, 0.9, 1.2, 2, 3.2, and 4 (or 5) s. We excluded the trials with delay lengths of 4 or 5 s consistent time conditions across mice (see below for more details). The probability of delay lengths followed the cumulative distribution function of exponential distributions (τ = 1.4 s) with a 0.2 s offset (Inagaki et al., 2019). This keeps the hazard function of the exponential distribution constant, disabling the mice from predicting when the go cue would be delivered (Zariwala et al., 2013). Trials in which the mice did not lick within 1.5 s after the go cue were classified as no-response trials. We excluded these no-response trials from the analysis. On average, each mouse performed 4.60±3.36 sessions over multiple days, with each session consisting of 121.43±7.57 right-lick trials and 123.30±27.84 left-lock trials on average (MEAN±STD).

**Figure 1.**
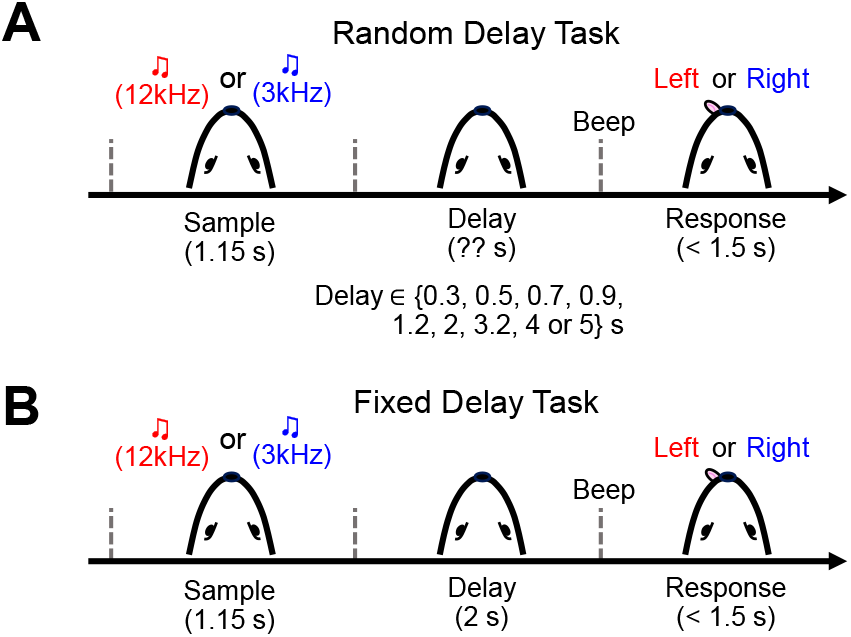
Task schematic for two types of auditory delayed-response tasks. (**A**) Task schematic for the random-delay task. As the trial starts, 3 or 12 kHz auditory cue is given for 1.15 s (Sample period). After a temporal delay (Delay period), mice lick right or left based on the auditory cue given during sample period (Response period). Delay length is randomly selected among eight delay lengths across trials. (**B**) In the fixed-delay task, the task structure is the same as that of the random-delay task, but delay length is fixed as 2 s across trials.

Separately, six mice were trained to perform the fixed-delay task, in which the delay length was fixed at 2 s, allowing the mice to predict go timing at the start of each trial. Each mouse performed 3.33 ± 1.32 sessions over multiple days, with each session consisting of 124.95±30.10 right-lick trials and 117.55±27.21 left-lick trials on average.

### Extracellular action potential analysis

Action potentials (spikes) were simultaneously recorded in left ALM using 64-channel Janelia silicon probes. Extracellular traces were recorded from left ALM and were band-pass filtered (300 – 6,000 Hz). Spike width was calculated as trough-to-peak interval in the averaged spike waveform. In the random-delay task, 841 of 1,005 recorded neural units were stably recorded through whole trials. Units with a width narrower than 0.35 ms were classified as putative fast-spiking neurons, whereas units with a width wider than 0.5 ms were defined as putative pyramidal neurons. Among total of 841 units recorded during the random-delay task, 106 units were putative fast-spiking neurons, and 718 units were putative pyramidal neurons. Other units with intermediate spike widths (0.35-0.50 ms, 17 out of 841) that could not be classified into either of these cell types were excluded from the analyses. As a result, 35.83±9.80 units (MEAN±SD) per session were used for analysis. In the fixed-delay task, 594 of 755 recorded neural units were stably recorded through whole trials. 62 units were putative fast-spiking neurons, 521 were putative pyramidal neurons, and 11 were excluded in the fixed-delay task. Thus, 29.15±16.14 units per session were used for analysis. Firing rates were computed with a 100 ms-sized squared bin at every 1 ms.

### Similarity between firing rate vectors at different time points

If population activities show persistent dynamics, activity patterns at different time points will be similar. On the other hand, if the dynamics of population activities fluctuate, activity patterns at close time points will be similar but those at distant time points will be different. Based on this idea, we analyzed how the firing rates of a population varied over time by measuring similarity between the firing rates of ALM populations at different time points.

In the random-delay task, we selected two set of trials with two different delay lengths (e.g., 2 s and 3.2 s). The delay lengths of 2 and 3.2 s were the longest durations used in the random-delay task. Therefore, it was suitable to examine persistent activity across delay period. Additionally, selecting different delay lengths allowed us to gather more trials, reducing the potential noise effect when averaging firing rates. For each set, we averaged the firing rate vector for N neurons across trials, at each time point. Each of these averaged firing rate vectors was then normalized to have unit length. We calculated the dot-product (i.e. similarity value) between the trial average vectors of every possible time point pair, then averaged those across sessions. In the fixed-delay task, the similarity was calculated as in the random-delay task, except that we randomly divided the trials into two sets per session.

### Selectivity

We first normalized firing rate using the baseline mean and standard deviation as follows:

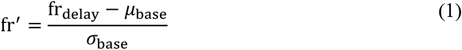

where fr_delay_ is a firing rate within a time window (50∼100 ms before delay ends) averaged across trials. *μ*_base_ and *σ*_base_ correspond to averaged baseline firing rate across trials and time (550∼50 ms before trial starts) and the standard deviation of that, respectively. We calculated selectivity as follows:

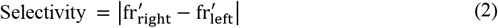

where 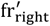 is the average of fr^′^across lick-right trials and 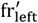 for lick-left trials.

### Alignment Index (AI)

We quantified how well the firing rates of neural populations were aligned on an arbitrary unit vector **V** using the Alignment Index (AI) (Elsayed et al., 2016), which is given as:

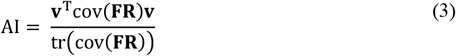

where cov(**FR)** is a covariance matrix of the firing rate data sized by N ×T.TR (N: number of neural units, T: time, TR: number of trials), tr(∙) is the trace operator, and **V** is an unit vector sized by N ×1. If **V** is a projection vector which maps N-dimensional firing activity onto a subspace, then AI represents the variance of firing rates captured by the subspace. AI is 1 if the covariance of the population activity is fully explained by the subspace, and 0 if the subspace explains no covariance of the population activity. For succinctness, we denote AI(Q, *S*) as AI during a task period Q (e.g., delay period) or specific time point (e.g., delay end), on subspace *S* (see below for more details of subspaces).

### Putative motor preparation and execution subspaces

Non-human primate studies have shown that neural trajectories of motor cortical neurons lie within different subspaces for different movement phases. With previous studies confirming that ALM populations show coordinated activities throughout movement preparation and execution (Elsayed et al., 2016; Lara et al., 2018), subspaces representing those coordinated activities can be different between movement preparation and execution (Economo et al., 2018; Inagaki et al., 2022a). Here, we called a subspace representing ALM population activities during movement preparation as a preparation subspace (***P***), and that during movement execution as a movement subspace (*M*). Preparation-specific activity and movement-specific activity are defined as neural activity projected on the preparation and movement subspaces, respectively. Preparation-specific activity is a latent component of neural population related to movement preparation during delay period, predicting upcoming movement information without making overt movement. Movement-specific activity is a similar latent component directly linked to movement execution.

We assumed that subspaces ***P*** and *M* should meet three requirements based on findings from other studies. First, preparation-specific activity at delay end (the preparatory end-state) should at least partially predict movement-specific activity right before making movement (Fig. 4A), as the preparatory end-state was an initial condition of subsequent movement. Second, ***P*** and *M* should be near-orthogonal to each other so that preparation-specific activity precludes muscle activity (Fig. 4A) (Kaufman et al., 2014; Elsayed et al., 2016; Stavisky et al., 2017; Kao and Hennequin, 2018). Note that the second requirement does not violate the first requirement: Trial-by-trial correlations can exist between the preparatory end-state and movement-specific activity right before making movement even when ***P*** and *M* are orthogonal to each other (Fig. 4A). Third, ***P*** and *M* should capture large variance of the population activity during movement preparation and execution, respectively. This is necessary because accurate licking is a goal of the task, thus neural components for preparing and executing licking should be prominently manifested in the population activity.

To estimate preparation-, and movement-specific activity, we inferred the projection vectors **P** and **m**, which linked high-dimensional population activities to low-dimensional activities on ***P*** and *M* respectively. We designed an objective function for learning **P** and **m** as follows:

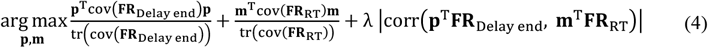

**FR**_Delay end_ indicates the firing rate matrix before delay end (averaged across 50∼80 ms before delay end), sized by N ×TR (N: number of neurons, TR: number of trials). **FR**_**RT**_ is a samesized firing rate matrix averaged over 20∼50 ms before the first lick. The first and second terms in eq. (4) allow **P** and **m** to capture the maximum variance of firing rates before the end of delay and prior to the first lick, respectively. The third term maximizes trial-by-trial correlations between the preparatory end-state and movement-specific activity before the first lick, implementing that the preparatory end-state predicts subsequent neural dynamics. The first and second terms in eq. (4) are bounded between 0 and 1. Similarly, the absolute value of the correlation term in eq. (4) falls within the range of 0 to 1. Therefore, we modified the objective function in eq. (4) as follows:

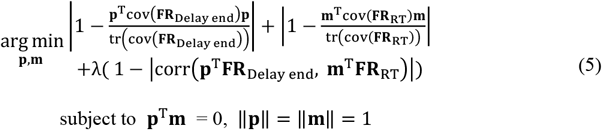

The optimization process could be accelerated by this modification. Even if **P** and **m** are orthogonal to each other, some neural variance during delay period could be captured by *M*. This is because dynamics that are not directly related to movement (e.g., non-selective ramping activities) may occur during the delay period (Yang et al., 2022). These non-movement components should not make activity on *M*. Therefore, we employed an optimization method with constraint to ensure that AI(Delay end, *M*) remains below 0.1, resulting in a subspace *M* that captures less than 10% of the neural variance during delay period. We learned **P** and **m** iteratively for each task using the MATLAB function *fmincon()*. At the end of each iteration, we performed Gram-Schmidt orthonormalization between **P** and **m** to secure orthogonality. The learning stopped when the first-order optimality measure was less than 10^−6^, while AI(Delay end, *M*) was less than 0.1.

A parameter λ balances between the third term and the first two terms. A larger λ puts more weight on increasing correlation during the optimization process, but this could hinder maximizing the first two terms. We empirically determined a minimum λ such that the resulting correlation after learning in the third term was significant for every session (*p* < 0.05). In this study, λ was set to 0.76 in the random-delay task and to 0.8 in the fixed-delay task.

To initialize **P** in the random-delay task, we conducted a principal component analysis (PCA) on trial-averaged firing activity matrix **FR** ∈ ℝ^N^ ^×T.C.D^ (N: number of neurons, T: time (150 ms after sample onset ∼ 50 ms before delay end), C: number of conditions (lick-right or lick-left), and D: number of delay lengths). Among the three PCs associated with the largest eigenvalues, we calculated the difference in PC activity between the lick directions. Then, we chose the PC that exhibited the highest degree of discrimination in distinguishing between licking directions as an initial **P**. Similarly, we conducted PCA on a trial-averaged firing activity matrix 0∼700 ms after making the first lick. Among the top three PCs, we selected one that captured the least variance of firing rates during delay period, as movement-specific activity is assumed to be absent during movement preparation. Then, we used the corresponding eigenvector as an initial **m**. In the fixed-delay task, the procedure was the same except that a trial-averaged firing activity matrix was sized by N ×T.C (N: number of neurons, T: time (50∼450 ms before go cue), C: conditions (lick-right or lick-left)).

After maximizing the objective function, we utilized the learned **P** and **m** to infer preparation-specific activity, 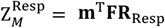 and movement-specific activity, 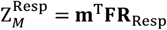 (**FR**_Delay_: firing activity during delay period, **FR**_Rep_: firing activity after the first lick). Note that we used averaged firing rates across trials in calculating AI on ***P*** and *M* in Figs. 4E-F.

### Random alignment index

We estimated how much task-irrelevant neural variance could be accounted for by the subspaces. We calculated this chance alignment, AI_rand_, using randomly generated projection vectors, **V**_rand_ (Elsayed et al., 2016; Jiang et al., 2020). We generated **V**_rand_ from multivariate normal distribution (mean vector: **0**, covariance: identity matrix). We calculated AI_rand_ as follows:

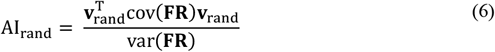

where cov(**FR)** is a covariance matrix of firing rate matrix **FR** ∈ ℝ^N^ ^×T.TR^ (N: number of neurons, T: time, TR: number of trials). We repeated this process 10,000 times. Then, we compared AI_rand_ with AI(Delay, ***P***) and AI(Response, *M*) in every session.

### Distance between the neural states on *P*-*M* dimension

Assuming that ***P*** and *M* could represent movement-related information, we inspected whether preparation-specific activity and movement-specific activity could predict RT on a single-trial basis. We reasoned that preparation- and movement-specific activities at delay end with the shortest RT across trials would be an optimal neural state to generate movement rapidly. Thus, we assumed that RT would increase as neural activities on ***P*** and *M* deviate more from this optimal neural state. We defined 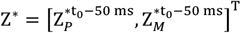 as a 2D vector of preparation-specific activity and movement-specific activities at 50 ms before delay end (t_0_-50 ms) that were obtained in a trial with the shortest RT, from a given session. The 50 ms offset prior to go cue ensures the exclusion of post-go-cue neural activity from the firing rate calculation, as the window size was 100 ms. We measured the Euclidean distance between Z^∗^ and Z(t_0_ − 50 **ms)** of the remaining trials as follows:

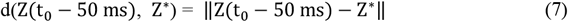

where 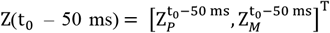 After calculating d(Z(t 50 ms), Z^∗^) for every trial, we calculated its correlation with deviations in RT (i.e., corr(d, **RT** − **min**(**RT)**)).

### Selectivity captured by the subspace

To estimate how much selectivity was captured by ***P*** and *M*, we first calculated firing rate matrix, **FR**_right_ (or **FR**_left_) averaged across lick-right trials (or lick-left). The matrices were sized by N ×T (N: number of neurons, T: time). We conducted singular value decomposition (SVD) on firing rate differences, **FR**_right_ − **FR**_left_. Using the set of right singular vectors **V**, we calculated total variance by projecting firing rate difference on singular vectors as follows:

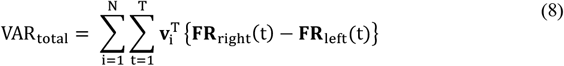

Similarly, we calculated neural variance on ***P*** by projecting firing rate difference on ***P***.

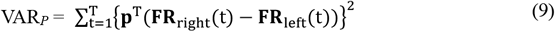

Total selectivity captured by ***P*** during delay period was given as:

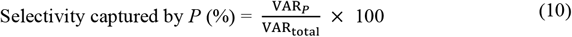

Substituting the projection vector in eq. (9) to **m**, the same calculations applied to calculate selectivity captured by *M*.

### Reconstruction of firing rates from preparation- and movement-specific activity

Assuming that the population activity can be decomposed into neural components of movement preparation (**FR**_*p*_) and execution (**FR**_*M*_), total firing rate can be expressed as

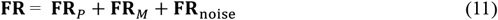

We reconstructed **FR**_*p*_ = **d**_*p*_**P**^T^**FR** and **FR**_*M*_ = **d**_*M*_**m**^T^ **FR** using decoder vectors **d**_*p*_ and **d**_*M*_ sized by N×1 (N: Number of neurons). The decoder vectors were estimated by minimizing the following objective function:

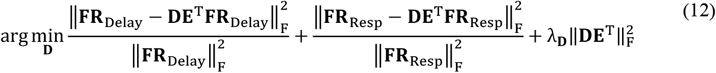

Where **D** = [**d**_*p*_ **d**_*M*_], **E** = [**P m**] and **‖**∙**‖**_F_ is the Frobenius norm of a matrix 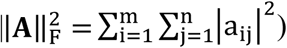 To decompose firing rates into unique components **FR**_*p*_ and **FR**_*M*_, we made **d**_*p*_ and **d**_*M*_ orthonormal to each other (Kobak et al., 2016). The reconstruction error is indicated by the first two terms of eq. (12), requiring 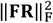 for normalization, while 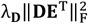 serves as a regularization term to prevent overfitting. We randomly sampled an 80% subset from total trials to minimize eq. (12), then selected λ_**D**_ that minimized the reconstruction error of the remaining 20%. To validate the learned decoder vectors **D**, we applied PCA to **FR**_Delay_ and **FR**_Rep_ using the same 80% subset. We extracted eigenvectors corresponding to the largest eigenvalue. Then, we reconstructed **FR**_Delay_ and **FR**_Rep_ using these eigenvectors and calculated reconstruction error for the test trials. We confirmed that there was no significant difference between reconstruction error using **D** and that using the eigenvectors (paired *t-*test, *p* = 0.899 in the random-delay task, *p* = 0.693 in the fixed-delay task). This verified that original **FR** could be sufficiently represented by reconstructed **FR** from preparation- and movement-specific activities.

### Distance between firing rates at different time points

We estimated how much firing activity at delay end, which generates no movement, would have to change to another one that can generate movement. To this end, we measured distance between firing rates at 50 ms before delay end (t_0_ − 50 **ms**) and at 50 ms before RT (t_**RT**_ − 50 **ms**). We randomly selected 20% of total trials and calculated their average firing rate 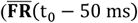 and 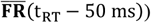 We decomposed these firing rates into movement preparation component, 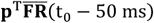 and execution component, 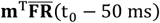

Based on the assumption that preparatory-specific activity becomes useless after go cue, we excluded preparation component 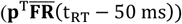 in measuring distance. We measured distance L between 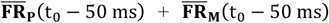 and 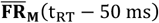 as follows:

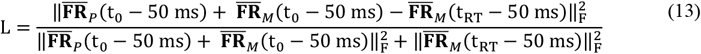

Where 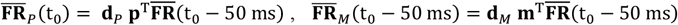 and 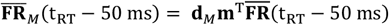 We repeatedly measured L by randomly sampling 20% of total trials 1,000 times. We averaged L across repetitions for each of right- and left-lick trials, then averaged the resulting two values.

## Results

### Behavioral outcomes and neural dynamics depend on the predictability of go timing

Aligning with other studies, in the random-delay task, ALM neurons’ firing activities rapidly increased then plateaued until the go cue (Fig. 2A) (Inagaki et al., 2019). Meanwhile, in the fixed-delay task, ALM neurons demonstrated gradually increasing activity throughout the delay period (Fig. 2B) (Inagaki et al., 2019).

**Figure 2.**
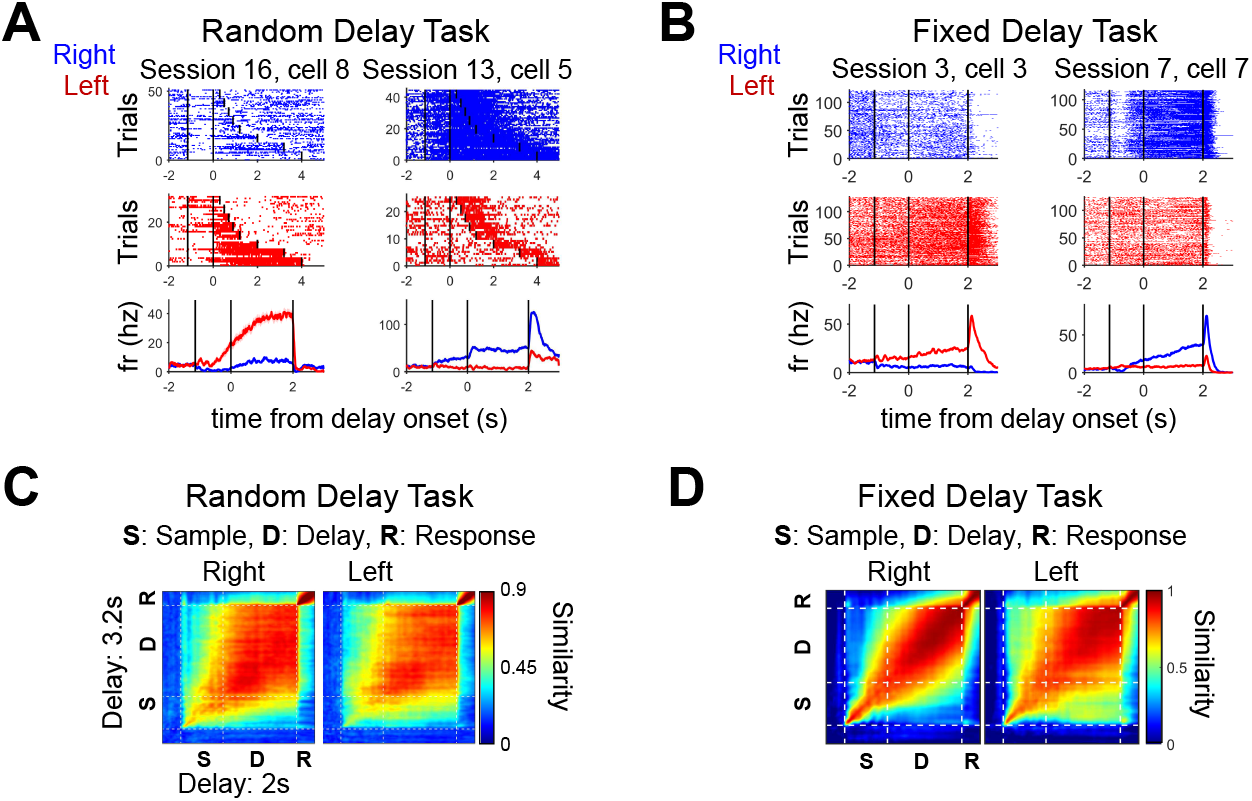
Different neural dynamics appeared depending on the predictability of go timing. (**A**) Example of neural responses during the random-delay task. Firing activities increased from sample period and persisted in the ready state in delay period. Vertical black lines indicate the start of Sample (−1.15 s), Delay (0 s), and Response period each. The top and middle are spike raster plots, and the bottom shows averaged firing rates across trials when 2 s delay was given. In spike raster plots, trials are randomly sampled for each delay length. Shading denotes SEM. (**B**) Example of neural responses during the fixed-delay task, similar to **(A)**. Unlike persistent dynamics in the fixed-delay task, neurons showed ramping dynamics during delay period. (**C**) Similarity between firing activities during 2 s (horizon) and 3.2 s (vertical) delay lengths in the random-delay task. The similarity was calculated at every time point and averaged across sessions. Dotted white lines separate each period (S: Sample, D: Delay, R: Response).(**D**) Similarity between firing activities in the fixed-delay task. The trials were randomly divided in half, and the similarity between the average firing rates of each set was calculated at every time point—these were averaged across sessions. Dotted white lines separate each period (S: Sample, D: Delay, R: Response).

To analyze these different activity patterns at the population level, we measured the similarities between firing rate vectors at every time point using dot-product (Figs. 2C-D). While similarity levels between adjacent time points was higher than that between more distant points in the fixed delay task (Fig. 2D), similarity was uniformly high during the whole delay period for the random-delay task (Fig. 2C) These similarity patterns support that different dynamics emerged at the population level depending on the predictability of go timing.

We also observed differences in behavioral outcomes. Rodents demonstrated higher decision accuracy and longer RT in the fixed-delay task than in the random-delay task (one-tailed *t-*test, *p* < 0.001) (Fig. 3A). To investigate the underlying neural mechanism of this difference, we calculated selectivity, indicating the degree of bias towards a particular licking direction in neuronal activity (right vs. left) (Li et al., 2015; Li et al., 2016a) (Fig. 3B). We expected that the degree of selectivity would be related to decision accuracy and elucidate the accuracy difference. Indeed, the degree of selectivity at the preparatory end-state was greater in the fixed-delay than in the random-delay task (one-tailed *t-*test, *p* < 0.001) (Fig. 3C). Further, the degree of selectivity at delay end increased as animals licked more accurately, only in the fixed delay task (**P**earson correlation coefficient, *r* = 0.716, *p* < 0.001) (Fig. 3D, right). There was no significant correlation between the degree of selectivity and behavioral accuracy in the randomdelay task (*p* > 0.1) (Fig. 3D, left).

**Figure 3.**
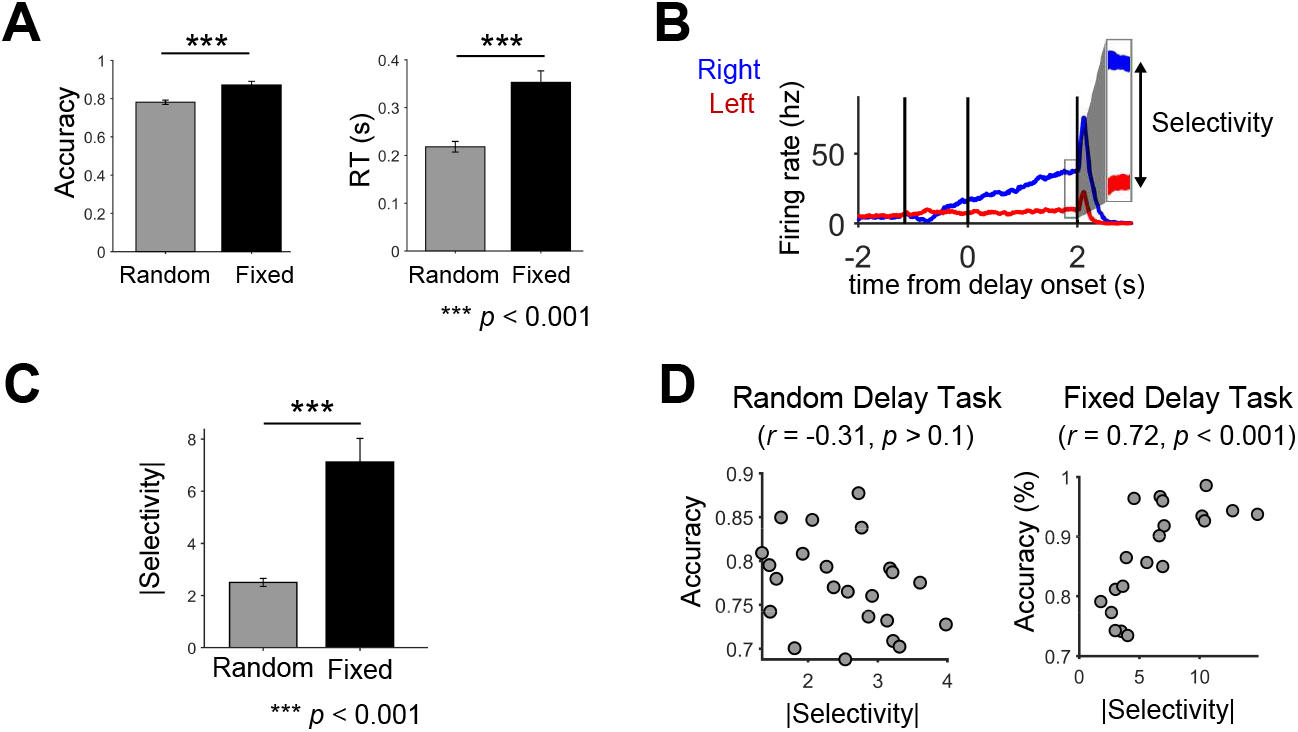
Behavioral outcomes and task-relevant neural characteristics are different between the tasks. (**A**) Behavioral accuracy and RT. Accuracy was higher and RT was longer during the fixed-delay task than those during the random-delay task (one-tailed *t-*test, *p* < 0.001). Error bar denotes SEM across sessions. (**B**) Selectivity at delay ended. Selectivity is firing rate difference between licking directions (right, left) normalized by baseline activity. (**C**) Averaged |Selectivity| after taking absolute on selectivity in (**B**). |Selectivity| in the fixed-delay task was significantly larger than |Selectivity| in the random-delay task (one-tailed *t-*test, *p* < 0.001). The error bar denotes SEM. (**D**) |Selectivity| in the random-delay task did not correlate with behavioral accuracy (*p* > 0.1) while |Selectivity| was positively correlated to behavioral accuracy in the fixed-delay task (*p* < 0.001).

**Figure 4.**
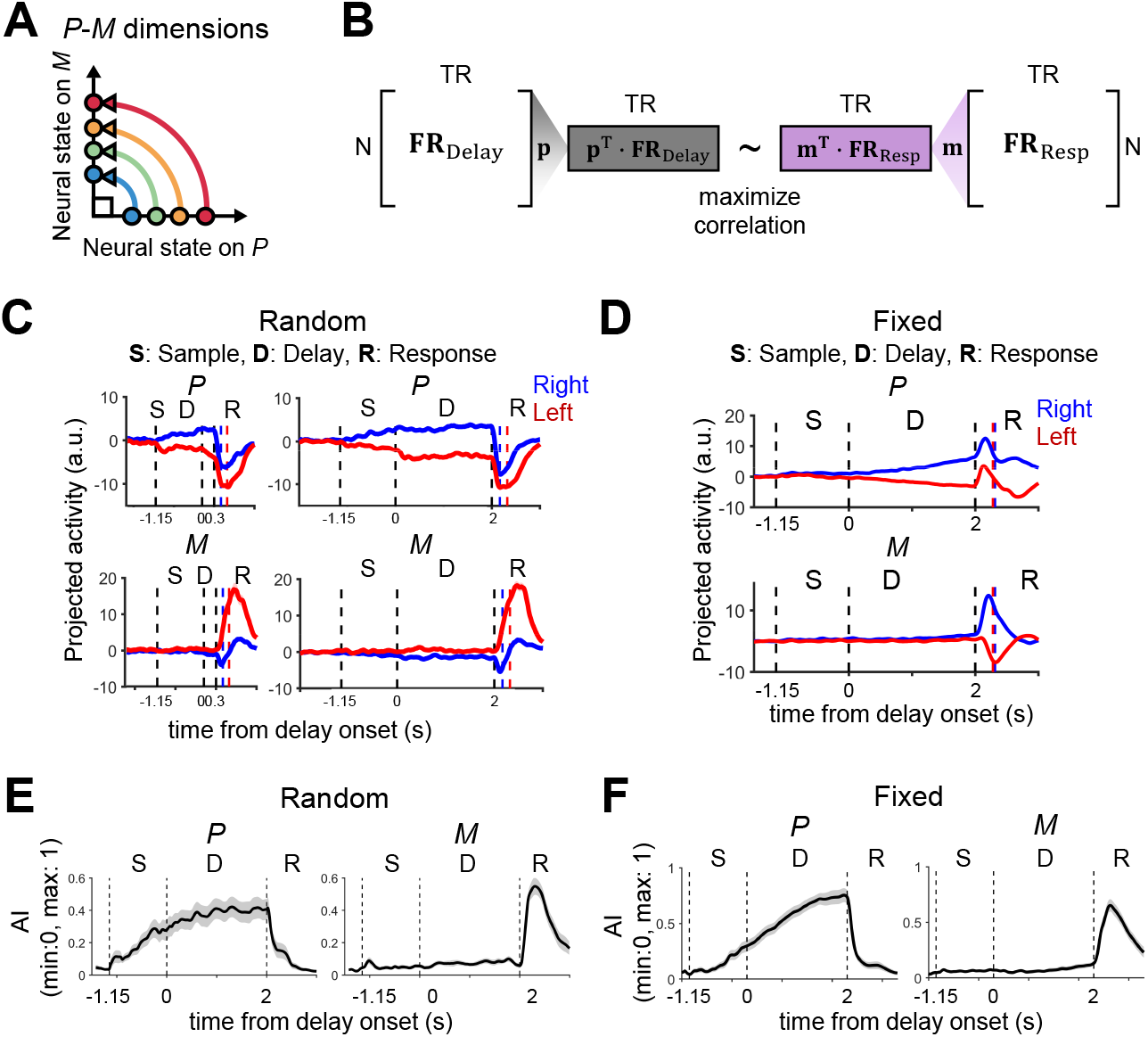
Population activities on inferred preparation and movement subspace. (**A**) The schematic diagram for correlated neural states of different movement phases. Each dot represents the neural state of a single trial. The neural state in preparation subspace ***P*** predicts the neural state in movement subspace *M*. ***P*** and *M* are orthogonal to each other so that neural activities on ***P*** do not show significant activities on *M*. Note that the neural state on ***P*** could be highly correlated to the neural state on *M* in trial-by-trial basis even ***P*** and *M* are orthogonal. (**B**) An algorithm seeks vectors **P** and **m** which are projection vectors to maximize the trial-by-trial correlation of the neural states during distinct epochs (N: number of neurons, TR: number of trials). (**C**) Example preparation- and movement-specific activities during the random-delay task (S: sample, D: delay, R: response). Left panels: 0.5 s delay, right panels: 2 s delay. Independent of delay lengths, the profile of preparation- and movement-specific activities appear similarly. Bold blue and red lines are projected activities averaged across trials and shaded background is standard error of the mean (SEM). Dotted blue and red lines indicate mean RT in lick-right and left each. (**D**) Example preparation- and movement-specific activities during the fixed-delay task (S: sample, D: delay, R: response). Dotted blue and red lines indicate mean RT in lick-right and left each (**E**) Alignment Index (AI) across delay period in the random-delay task (delay length: 2 s). AI on ***P*** increases over sample period and flattens when delay starts (left panel). Meanwhile, AI on *M* remains low during delay period and rises after go cue (right panel). Gray shadow represents SEM across sessions. (**F**) Alignment index (AI) averaged across sessions in the fixed-delay task. AI on ***P*** gradually increases until go cue (left panel). AI on *M* remains low during delay period and rapidly increases after go cue (right panel). Gray shadow represents SEM across sessions.

The dynamical system, being near-autonomous, assumes that an initial condition largely determines the neural dynamics for the subsequent move (Churchland et al., 2010; Remington et al., 2018; Wang et al., 2018; Kao et al., 2021). Thus, if the preparatory end-state for both tasks are similar, both transitions to the subsequent movement initiation should span similar time (i.e., similar RT) as well. But as shown in Fig. 3A, RT was significantly longer in the fixed-delay task, indicating that the preparatory end-state are in different forms for the two tasks.

Based on these neural and behavioral results, we hypothesize that different movement preparation strategies would be adopted by motor cortical neurons depending on the predictability of go timing. When go timing is unpredictable, preparatory activity quickly reaches a neural state that can produce a rapid licking movement and persists in that state until the go cue is given. However, it results in less accurate behavioral performance as evidenced by low selectivity (Fig. 3C). On the other hand, when go timing is predictable, neural activity shapes the preparatory end-state to increase behavioral accuracy as evidenced by high selectivity (Fig. 3C).

### ALM population activity is segregated in preparation and movement subspaces

To test our hypotheses, we extracted preparatory-, and movement-specific activities by inferring preparation and movement subspaces ***P*** and *M* (Methods). 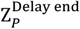 and 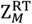 showed significant trial-by-trial correlations in every session. The preparatory end-state could predict a neural state of movement initiation, as the absolute value of **P**earson correlation coefficient *r* was 0.54±0.03 for the random-delay task and 0.67±0.03 for the fixed-delay task (*p* < 0.05 for every session in both tasks). Also, movement-specific activity was largely absent until the end of delay. It started to increase only right before the first lick occurred (Figs. 4C-F), similar to previous findings by Inagaki et al. (Inagaki et al., 2022b).

We calculated AI of every ms (100-ms bin width) to measure how much neural variance the inferred subspaces capture at each time point (Methods). In the random-delay task, both AI(Delay, ***P***) and AI(Delay, *M*) remained relatively persistent during delay period (Fig. 4E). This demonstrated that the dynamics of preparatory activity did not progress further towards preparation-or movement-specific activity during delay period, even when additional time was available (Fig. 4E). In the fixed-delay task, AI(Delay, ***P***) gradually increased during delay period (Fig. 4F, left), while AI(Delay, *M*) was largely absent as in the case of the random delay condition (random: 0.073±0.015, fixed: 0.066±0.014, MEAN±SD across delay period) (Figs. 4E-F).

We found that both AI(Delay, ***P***) and AI(Response, *M*) were significantly higher than AI_rand_ in every session (one-tailed *t-*test, *p* < 10^−9^ for every session), supporting that subspaces ***P*** and *M* captured a significant amount of the variance of population activity during different periods of the task.

### Movement-related information is encoded on preparation and movement subspaces

To verify if the inferred subspaces ***P*** and *M* were related to the preparation and execution of directional licking, we tested whether movement-related information such as RT and licking-direction were encoded on ***P*** and *M*. We first examined whether RT could be predicted by preparation-, and movement-specific activities. We found that **RT** − **min**(**RT)** was correlated with the Euclidean distance from Z to Z^∗^ across trials (Methods). The resulting correlation coefficients were significantly larger than 0 in both tasks (one-tailed t-test, p < 0.05), which indicated that the farther Z deviated from Z^∗^, RT increased (Fig. 5A).

**Figure 5.**
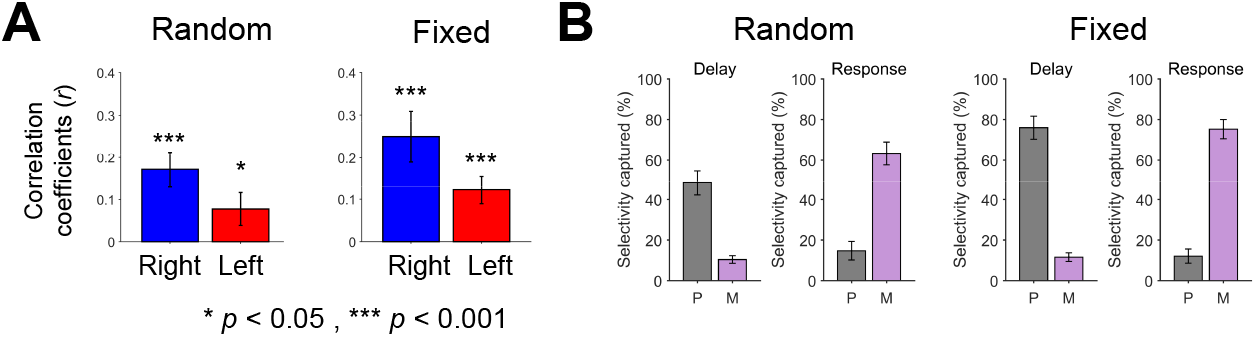
The neural state on ***P***-*M* dimensions predicts behavior in a single trial basis. (**A**) Distance between Z and Z^∗^ is positively correlated to **RT** − **min**(**RT)** across trials. Correlation coefficient is significantly larger than 0 (one-tailed paired *t-*test). Error bar indicates SEM across sessions. (**B**) Selectivity captured by ***P*** and *M*. Error bar indicates SEM across sessions.

Next, we quantified how much directional selectivity was captured by ***P*** and *M* (Methods). We found that a significant amount of selectivity was captured by both ***P*** and *M* (Fig. 5B). In the random-delay task, 58.90 ± 6.61% of the population selectivity during delay period was captured by ***P*** and *M* (48.46±6.14% for ***P***, 10.44±1.91% for *M*) and 78.00±4.28% during response period (14.78±3.08% for ***P***, 63.22±13.18% for *M*) (MEAN±SEM across sessions). In the fixed-delay task, 87.54±4.99% of the population selectivity during delay period was captured by ***P*** and *M* (75.93±5.67% for ***P***, 11.61±2.10% for *M*) and 87.34±3.47% during response period (12.08±2.70% for ***P***, 75.26±16.83% for *M*) (MEAN±SEM across sessions). The previous study by Li et al. reported that neural manifolds that were inferred to discriminate licking direction captured 65.6±5.1% of the population selectivity while mice performed a similar perceptual decision task (Li et al., 2016b). Thus, the captured selectivity in the present study were on par with that in the previous study. Collectively, we verified that neural activities on ***P*** and *M* could encode movement information such as RT and licking directions.

### Persistent activity during the random-delay task rapidly responds to the go cue under temporal uncertainty

We hypothesized that, in the random-delay task, neural dynamics would quickly reach a neural state ready to initiate licking movement anytime, and persist until the go cue (persistent dynamics, Figs. 2A, C). To test our hypothesis, we estimated when neural activity became ready to make a licking movement during delay period. Especially, we examined when movement-related information such as RT could be predicted from preparation-specific activity.

Lara and colleagues reported that preparation-specific activity increased and peaked several ms before movement initiation. This incremental occupancy of preparation-specific activity among total neural activity indicated that preparation-specific activity became more prevalent as the population activity approached the preparatory end-state (Lara et al., 2018). Thus, we used this occupancy to estimate the extent to which the population activity was ready to initiate a movement. We assumed that high occupancy of preparation-specific activity predicts shorter RT. We used AI(Delay, ***P***) to measure occupancy. Furthermore, to investigate whether movement-specific activity during delay period reaches the desired state faster, leading to quicker movements after go cue, we tested if AI(Delay, *M*) was negatively correlated with RT. Hence, if neural activity at a given time point during delay period is ready to move, its AI on ***P*** and *M* would show negative correlations with RT.

We calculated correlations between AI and behavioral outcomes (RT, accuracy) across sessions. Here, we conducted a correlation analysis across sessions because decision accuracy and AI were calculated once in each session. In the random-delay task, we found that decision accuracy was not correlated with AI(Delay, ***P***) nor AI(Delay, *M*) (Fig. 6A). However, negative correlations between AI(Delay, ***P***) and RT were observed before delay started, until the end of delay (Fig. 6B, top). This early appearance of a negative correlation also existed between AI(Delay, *M*) and RT (Fig. 6B, bottom). We measured a time lag from trial onset to the first emergence of a significant negative correlation. The first significant correlation appeared even during sample period (Fig. 6C). Although the correlation coefficient was slightly jittered during sample period, it became stable when delay started (Fig. 6B). Also, we observed that the average correlation between the first appearance of negative correlation to go cue was negative regardless of delay lengths (Fig. 6D).

**Figure 6.**
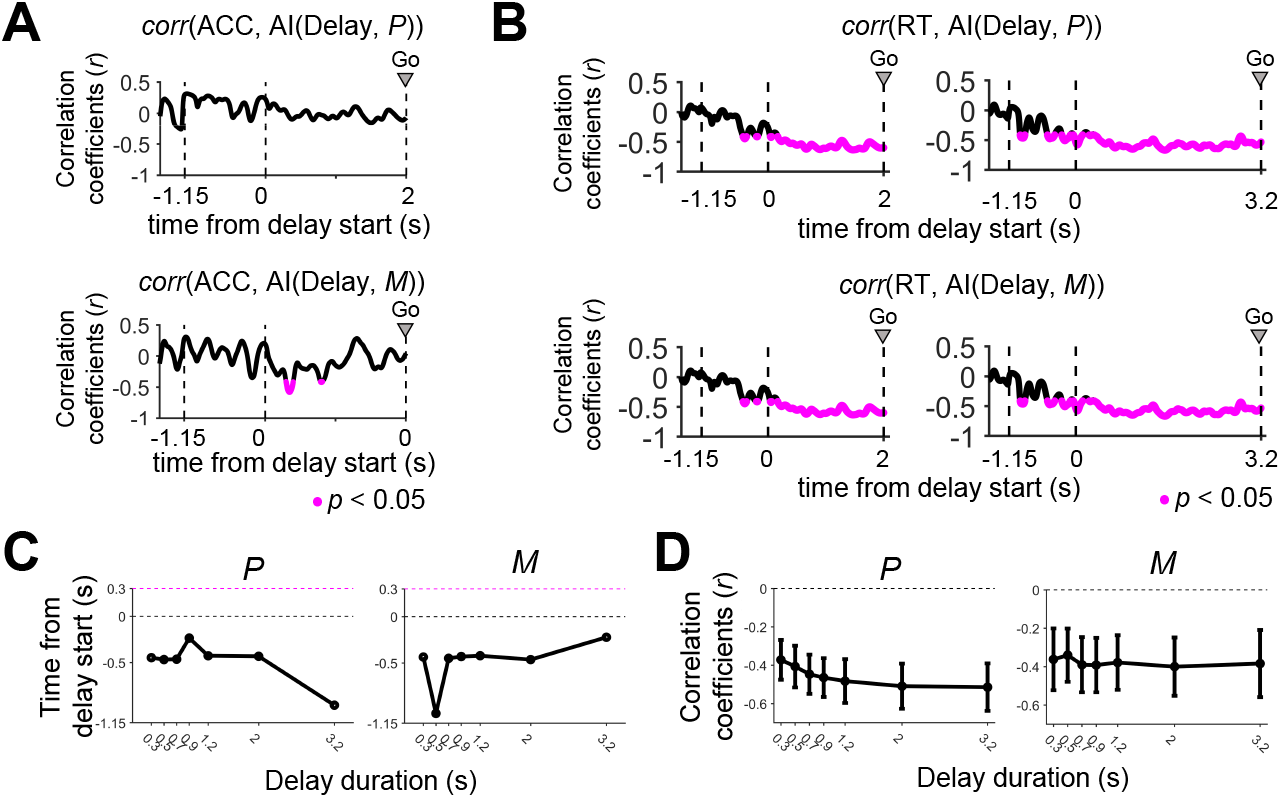
Preparation- and movement-specific activity appears right after delay starts. (**A**) Example correlation between AI and decision accuracy (delay length: 2 s). S: Sample, D: Delay. (**B**) Example correlation between AI and RT (left panel: 2 s delay, right panel: 3.2 s delay). The magenta dot represents the time point at which correlation is significant (*p* < 0.05). Significant and negative correlation between RT and AI appears before delay period. (**C**) The first significant and negative correlation between AI and RT appears before delay begins regardless of delay length. Black dotted line indicates time point when delay begins (0s). Magenta color dotted line indicates the shortest delay length given in the random-delay task. (**D**) Average correlation coefficient between AI and RT from the time when the first significant and negative correlation appears to the moment go cue is presented (MEAN±SEM). Dotted line indicates zero correlation coefficient.

Previous studies reported that RT becomes shorter when more time is given to prepare movement with predictable go timing, which is referred to as “RT savings” (Fig. 7A) (Riehle et al., 1994; Crammond and Kalaska, 2000; Churchland et al., 2006b; Churchland and Shenoy, 2007). Accordingly, we tested if RT decreased as delay length increased in the random-delay task. However, statistical analyses revealed that such RT savings were not present in the random-delay task (ANOVA, *p* > 0.1 for testing the effect of delay length on RT; linear regression *p* > 0.1) (Fig. 7B). Moreover, there was no significant difference in AI(Delay end, ***P***) among different delay lengths (ANOVA, *p* > 0.1 for testing the effect of delay length on AI(Delay end, ***P***); linear regression *p* > 0.1) (Fig. 7B). This indicated that the absence of an inverse relationship between RT and delay lengths was related to the lack of change in AI(Delay, ***P***) at delay end. To explore why AI(Delay end, ***P***) did not increase with delay length, we examined how AI(Delay, ***P***) evolved from trial onset to delay end. We observed that AI(Delay, ***P***) did not reach the maximum value at the end of delay when delay was short (e.g., 0.3 s), and decreased after reaching the maximum when delay was long (3.2 s) (Fig. 7C). Yet, we noticed RT variation over delay lengths in a smaller scale (the shortest RT was 0.215 s at a 1.2 s delay, and the longest RT was 0.226 at 3.2 s delay on average, Fig. 7B). We then found an negative correlation between AI(Delay end, ***P***) and RT (Fig. 7D) (Pearson correlation coefficient, *r* = - 0.799, *p* = 0.031). Thus, AI(Delay end, ***P***) – which represents how close preparatory activity is to the preparatory end-state – still elucidated variation of RT even in the absence of RT savings.

**Figure 7.**
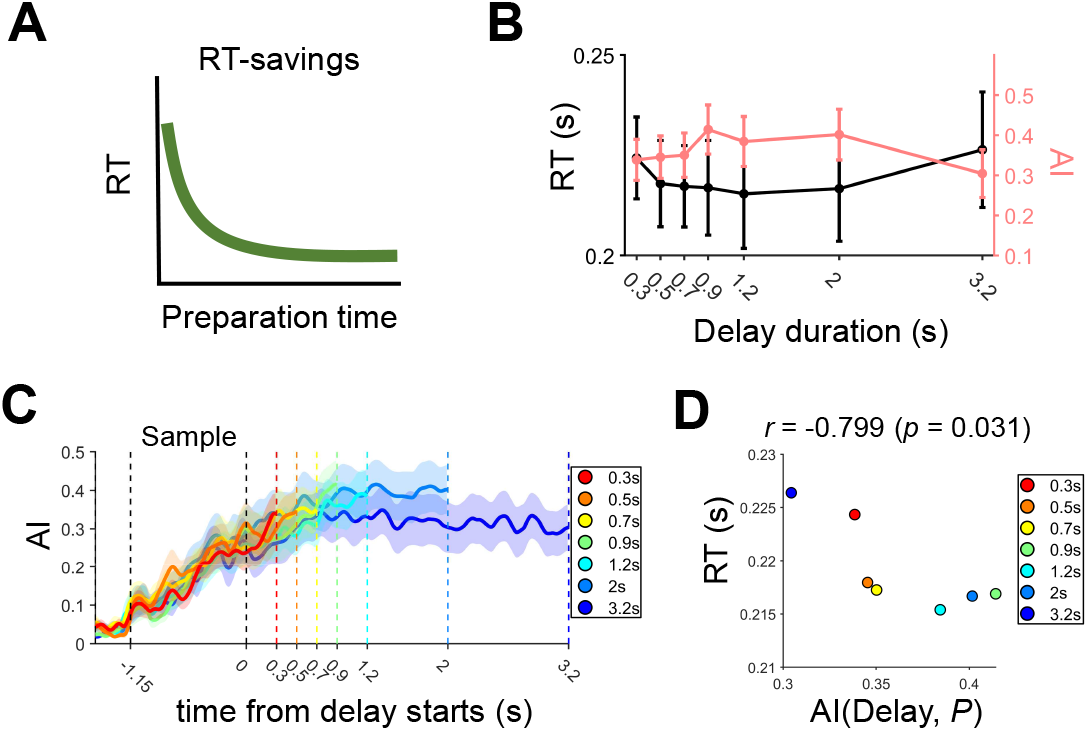
RT-savings earned by preparation time is removed in the random-delay task. (**A**) An illustration of the RT savings where RT is decreased with the duration of movement preparation time. (**B**) RT across delay lengths. There was no significant difference in RTs between delay lengths (ANOVA, *p* > 0.1; linear regression, *p* > 0.1 for slope). Error bar indicates SEM across sessions. Also, there was no significant different in AI between delay lengths (ANOVA, *p* > 0.1; linear regression, *p* > 0.1 for slope). (**C**) AI on ***P*** in the random-delay task. ALM activities were mostly persistent, but AI on ***P*** slightly increased during early phase of delay. If delay length becomes longer, AI on ***P*** decreased slightly, but this change was not statistically significant (ANOVA, *p* > 0.1). Shading represents SEM across sessions. (**D**) At the end of delay, AI on ***P*** and RT are inversely related (**P**earson correlation coefficients, *r* = -0.799, *p* = 0.031).

### The preparatory end-state maximizes in accuracy but accompanies long RT in the fixed-delay task

Behavioral accuracy was higher in the fixed-delay task than in the random-delay task (Fig. 3A). Based on this result, we hypothesized that neural dynamics would shape the preparatory endstate in a way that makes the following movement more accurate when the go cue is predictable. To test this, we measured correlations between AI and accuracy in the fixed-delay task: unlike in the random-delay task, AI(Delay, ***P***) gradually increased during delay period (Fig. 4F, left) and was positively correlated with accuracy (Fig. 8A, top). This showed that when preparation-specific activity occupied more of the population activity, decision accuracy was higher. It also indicated that the preparatory end-state, which was used to construct ***P***, represents neural information related to motor decision on licking direction. AI(Delay, *M*) showed no correlation with accuracy (Fig. 8A bottom).

**Figure 8.**
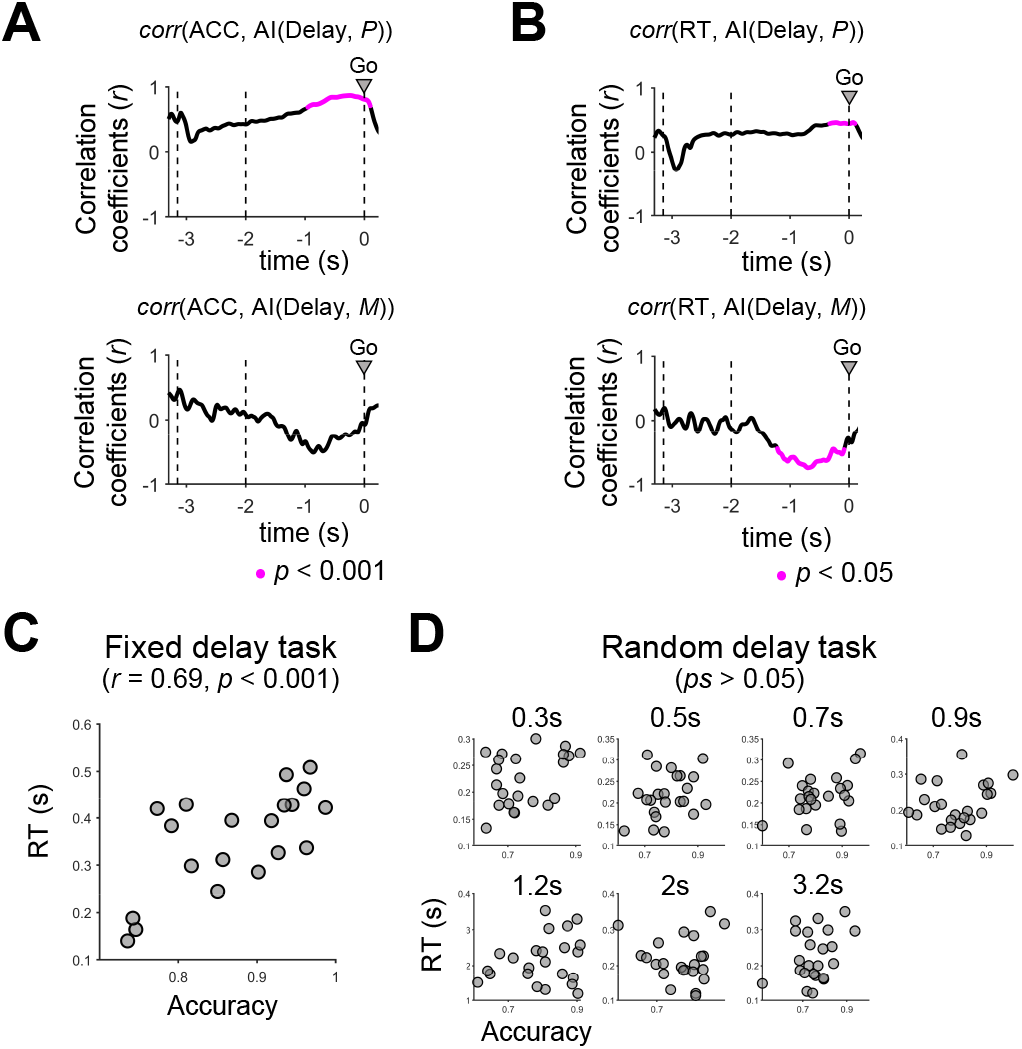
Preparation- and movement-specific activity encode behavioral outcomes in the fixed-delay task. (**A**) Correlation between AI and accuracy. The magenta-colored dot indicates the time point when correlation is significant (*p* < 0.001). A significant positive correlation between accuracy and AI(Delay, ***P***) appears right after delay starts (top). Movement-specific activity does not correlate with accuracy (bottom).(**B**) Correlation between AI and RT. RT shows a significant positive correlation with AI(Delay, ***P***) at late delay, which implies that dominant preparation-specific activity results in longer RT (top). Similar to the random-delay task, high AI(Delay, *M*) results in shorter RT but a significant and negative correlation starts to emerge from -1.22 s in the fixed-delay task, which is delayed than the random-delay task (bottom). (**C**) Behavioral accuracy is positively correlated with the RT in the fixed-delay task (*r* = 0.69, *p* < 0.001). (**D**) In the random-delay task, there was no significant correlation between behavioral accuracy and RT for every delay length (*ps* > 0.05).

We also observed that RT was slower in the fixed-delay task than in the random-delay task. Thereby, we measured correlations between AI and RT in the fixed-delay task and found a negatively correlation across sessions (Fig. 8B, bottom). This was similar to the random-delay task as expected, since AI(Delay, *M*) reflected how close the neural activities during delay and at movement initiation were. We observed that the time lag from delay onset to the first emergence of a significant negative correlation was longer in the fixed-delay (< 0.8 s after delay starts) than in the random-delay task (sample period). This indicates that during the delay period, neural dynamics gets ready to make movements more slowly in the fixed-delay task than in random-delay. AI(Delay, ***P***), on the other hand, was positively correlated with RT near the end of delay (Fig. 8B, top). The positive correlations of AI(Delay, ***P***) with both RT and accuracy indicated that the preparation subspace ***P*** would be altered in the fixed-delay task rather than in the random-delay task, even though ***P*** was learned in an unsupervised manner without using the licking behavior information.

In fact, behavioral data revealed that accuracy and RT were positively correlated with each other in the fixed-delay task (**P**earson correlation coefficients, *r* = 0.69, *p* < 0.001) (Fig. 8C), but not in the random-delay task (*ps* > 0.05 for every delay length) (Fig. 8D). Taking this with the behavioral observations, we postulated that the preparatory end-state was shaped towards greater behavioral accuracy but with longer RT, in the fixed-delay task.

We further examined why increasing behavioral accuracy accompanied longer RT. We defined 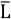 as the distance between the preparatory end-state and movement-specific neural state 50 ms before the first licking (Fig. 9A). We selected this timing of 50 ms prior to movement onset to take account of the time it takes for neural activity to translate into muscle activity. To calculate 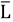, we reconstructed firing rate vectors from ***P*** and *M* to include only neural components of movement preparation and execution (Methods). Note that we did not measure distance in ***P***-*M* dimensions because a change in the neural state from preparation to movement could be different among neurons.

**Figure 9.**
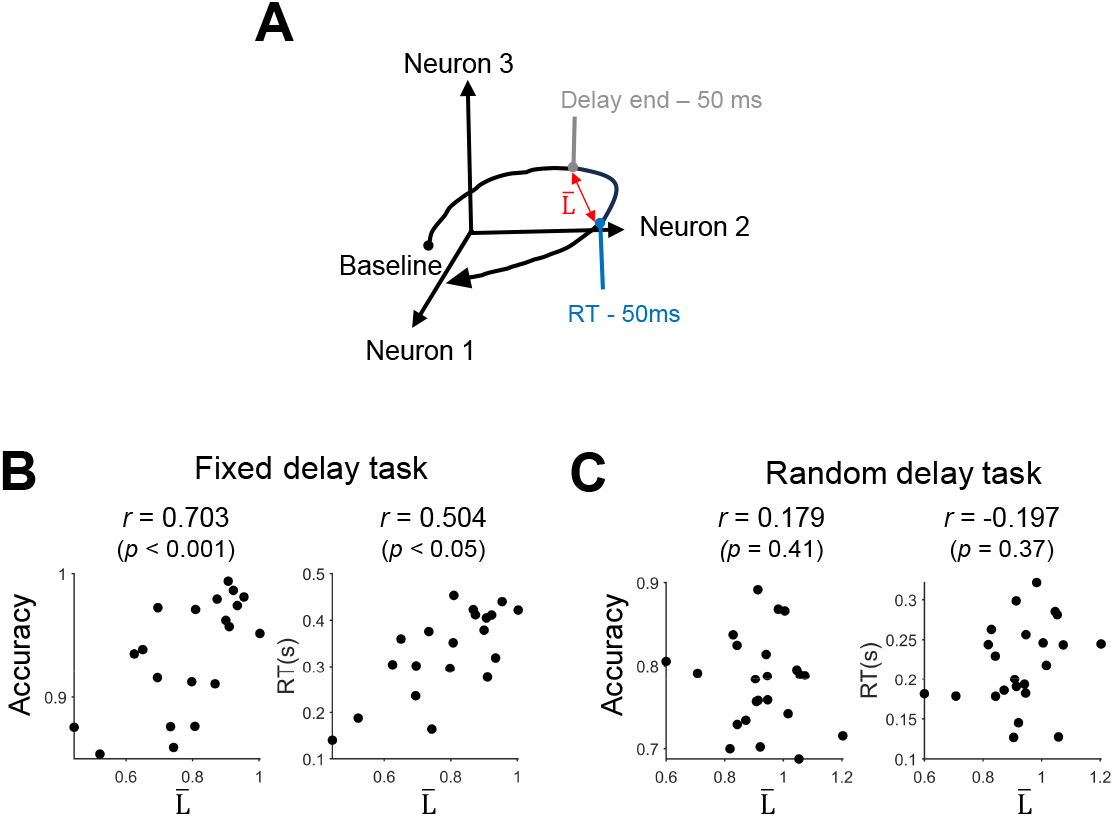
Accuracy increases as distance (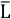) between the neural states at delay termination and movement initiation increases. (**A**) The schematic diagram of distance 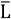 between the neural states. 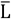 is the Euclidean distance between the neural states on 2D ***P***-*M* spaces. Here we measured the distance between preparation- and movement-specific activity at 50ms before delay end and 50 ms before movement initiation. (**B**) In the fixed-delay task, accuracy increased as 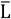 increased (**P**earson correlation coefficients, *r* = 0.6450 *p* = 0.0002). Also, RT increased as 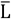 increased (**P**earson correlation coefficients, *r* = 0.7186, *p* = 0.0004). (**C**) In the random-delay task, 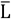 was positively correlated with RT but not significant.

If 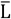 was positively correlated with accuracy, preparatory end-state would be formed in a direction of increasing accuracy as 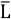 increased. However, it would also take longer to initiate movement because of the larger distance from the preparatory end-state to movement-specific neural state (i.e., longer RT as 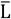 increases). Indeed, we found that both accuracy and RT increased as 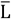 increased (Fig. 9B). We then controlled the effect of 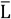 to investigate accuracy and RT’s dependency on 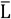. Controlling so in the fixed-delay task, there was no longer a speed-accuracy trade-off between accuracy and RT (linear partial correlation, *p* = 0.125). Note that correlations between 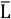 and RT, or between 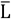 and accuracy, were not found in the random-delay task (Fig. 9C).

## Discussion

We demonstrated that preparatory activity is systemically altered depending on the predictability of go timing. We inferred preparation and movement subspaces from ALM population activity to decompose preparation-, and movement-specific activity (Figs. 4-8)). When go timing is unpredictable (e.g., random-delay length), neural dynamics quickly reached the preparatory end-state and maintained it until movement generation (Figs. 2, 6). On the other hand, neural dynamics reached the preparatory end-state gradually when there was no need to cope with temporal uncertainty (e.g., fixed-delay length), in order to more accurately distinguish between alternative licks (Figs. 2B, D). Behaviorally, however, predictable go timing resulted in higher accuracy and longer RT (Fig. 8C). This is because, as accuracy increased, the preparatory end-state moved further away from the neural state for the onset of movement, resulting in longer RT (Fig. 9B). Thus, motor cortical dynamics may evolve to generate an end-state for increased movement accuracy, but with no free lunch – the end-state also becomes more distant from movement-initiation states, resulting in longer RT in the delayed-response task with fixed delay length. Our findings suggest a speed-accuracy trade-off in which motor cortex (ALM) can flexibly generate preparatory activity using temporal resources to enhance behavioral accuracy at the cost of shifts in the preparatory state.

We find two main differences in the movement preparation process depending on the predictability of go timing: the speed at which the preparatory end-state is reached and the behavioral information that the preparatory end-state encodes. When go timing is unpredictable, neural activity rapidly reaches the end-state and persists there until cued to initiate movement (Figs. 2C-D). A relatively short time to collect sensory evidence may explain the low decision accuracy in the random-delay task. Meanwhile, if preparation duration is fixed, preparatory activity ramps up throughout the whole delay period. When the go cue is unexpectedly not delivered at the predicted go timing in the fixed-delay task, preparatory activity persists its current state, like in the random-delay task (Tanaka, 2007; Inagaki et al., 2022b).

In both tasks, behavioral responses became faster as neural activity progressed into movement subspace *M* during delay period (Figs. 6B bottom, 8B bottom). It supports the idea that the neural subspace representing movement execution (i.e., movement subspace *M*) would remain consistent between the tasks. However, what and how activities on the preparation subspace ***P*** encode for upcoming behavior were different between the tasks (Figs. 6, 8). In the random-delay task, greater occupancy of preparation-specific activity among population activity resulted in a shorter RT, while it did not predict decision accuracy. In the fixed-delay task, however, the greater occupancy resulted in higher accuracy but longer RT. A recent study confirmed the observation that the same preparation subspace activity appears when pursuing the same movement under different movement initiation conditions (e.g., reaching after a fixed-delay or self-cued reaching) (Lara et al., 2018). Therefore, whether the movement is cue-, or self-initiated, the same preparation subspace is represented. However, unlike our study, the movement preparation process in Lara et al. (2018)’s study did not involve decision-making, because the target was fully specified. Thus, we surmise that systematic changes in a neural network level would emerge through learning different goal-directed movement tasks. Also, these changes would engage cognitive processes to attain the goal, resulting in differential formations of neural dynamics of preparatory activity.

One potential source that may contribute to that differentiation is ramping input, which delivers task-relevant timing information (Kunimatsu et al., 2018). **P**revious studies showed that preparatory activity approaches one of the discrete attractors that each match with a different movement option (Shenoy et al., 2013; Inagaki et al., 2022a). When go timing is predictable, timing information is reflected in preparatory activity via a ramping input from other brain regions (Inagaki et al., 2018; Inagaki et al., 2019; Finkelstein et al., 2021; Inagaki et al., 2022a). The input drives the dynamics of preparatory activity to move the attractors apart, predisposing motor cortical neurons to be able to distinguish between movement options (Fig. 3C) (Finkelstein et al., 2021). However, the ramping input is absent when go timing is unpredictable (Inagaki et al., 2019). In practice, ALM neurons encoding ramping signals are significantly overlapped with those encoding choice of movement option (Yang et al., 2022). Collectively, the presence or absence of ramping input into ALM circuits would result in modifications of preparatory activity.

It has been assumed that one of the roles of movement preparation is to make rapid responses. For example, longer preparation time yields shorter RT (Rosenbaum, 1980; Riehle and Requin, 1989; Duan et al., 2021). Moreover, when the preparatory end-state is disrupted by electrical microstimulation, RT-savings gained through movement preparation are lost (Churchland and Shenoy, 2007). However, when go timing is unpredictable, we found long RT and persistent decision accuracy (ANOVA, *p* > 0.05 for delay length) even during longer delay thus more time to prepare for movement (Fig. 7B). The level of movement readiness did not increase as well (Fig. 7B). These results suggest that, when delay length is uncertain, neural dynamics do not proceed further toward the neural state of movement (i.e., neural response just before the first lick) thus yield constant RT across different delay lengths. Other studies have demonstrated that RT-savings appeared when delay length was randomly sampled from a uniform distribution (Duan et al., 2021). Yet, it is known that a hazard rate – probability that the go cue will be given in the remaining delay period – keeps increasing until the end of delay, in a uniform delay length distribution (see Zariwala et al. (2013) for more information). Since this allows prediction of go timing, preparatory activity would proceed towards the neural activity of movement as time passes, and RT would be shortened accordingly. On the contrary, in our data, delay length was randomly sampled from an exponential distribution, which is known to make a hazard rate constant over the whole delay period (Inagaki et al., 2019), so as to rule out a possibility of predicting go timing. Therefore, RT-savings may occur only when the probability of presenting the go cue increases as the waiting time increases (Janssen and Shadlen, 2005).

Speed-accuracy trade-off has been explained using a serial accumulate-to-threshold framework (Ratcliff and Smith, 2004; **P**almer et al., 2005; Heitz, 2014; Servant et al., 2019). According to this framework, decision-making is relatively inaccurate but fast when decisions are made by accumulating only a few pieces of sensory evidence quickly to reach a decision threshold. In contrast, decision-making is more accurate but slower when subjects accumulate more sensory evidence until the threshold. Similarly, multiple brain regions have been reported to show growing activity until threshold during decision-making. For example, neural activity in superior colliculus and the lateral intraparietal area (LI**P**) during saccadic response task accumulates evidence until the activity reaches a fixed threshold (Ratcliff et al., 2003). Also, premotor cortex shows an increase in baseline activity the motor task more rapidly reaches for threshold (Churchland et al., 2006a).

In the fixed-delay task, however, time given to collect sensory evidence was fixed. Also, temporal delay is interleaved between sample and response periods, thus what determined a balance between speed and accuracy would not be an accumulation of sensory evidence. We found that the transition from the preparatory end-state to the neural state of movement onset became distant as decision-accuracy was high. This could be because modifying neural activities to improve the decision-making process would affect movement preparation. For example, selectivity in ALM becomes salient and correlations between selective neurons are re-established through task learning (Komiyama et al., 2010). However, it is unclear why this modification should accompany un-optimal response speed. This could be because an intrinsic structure of a neural network before learning shaped the neural dynamics (Sadtler et al., 2014). Future studies on how the motor cortical neurons optimize task performance through learning will shed light on potential neural substrates of speed-accuracy tradeoff in motor cortex.

In the present study, we do not compare the kinematics of licking between the tasks. Thus, it is difficult to argue that kinematically identical licking can be produced from the different preparatory end-states. However, licking movement typically stays consistent, especially when mice skillfully learned the task (Bollu et al., 2021; Xu et al., 2022). Also, ALM is premotor cortex, thus ALM neurons encode intended lick angle and non-kinematic aspects of licking such as a licking initiation. Specific kinematics of licking such as tongue length and its rate are encoded in other regions of motor cortex such as the tongue and jaw regions of motor cortex (Xu et al., 2022). Thus, we speculate that preparation subspace is altered, because what and how preparation-specific activity encodes becomes different while pursuing the same action.

Our results help to provide a neural explanation for the task strategies that vary with our internalized time limits. Our study does not explain the optimality of the adopted task strategy in a given task structure. Future studies could shed light on how cognition shapes motor cortical dynamics and how, in turn, intrinsic constraints of motor cortical dynamics affect cognition.

## Acknowledgments

We thank the Svoboda laboratory for the contribution of data publicly at the FigShare, a datasharing website. This study was supported by the Brain Convergence Research **P**rograms of the National Research Foundation (NRF) funded by the Korean government (MSIT) (NRF 2021M3E5D2A01019542) and the Alchemist **P**roject Brain to X (B2X) **P**roject funded by the Ministry of Trade, Industry and Energy (No. 20012355; NTIS No. 1415181023).

## References

Afshar A, Santhanam G, Yu BM, Ryu SI, Sahani M, Shenoy KV (2011) Single-trial neural correlates of arm movement preparation. Neuron 71:555–564.

Bollu T, Ito BS, Whitehead SC, Kardon B, Redd J, Liu MH, Goldberg JH (2021) Cortex-dependent corrections as the tongue reaches for and misses targets. Nature 594:82–+.

Churchland MM, Shenoy KV (2007) Delay of movement caused by disruption of cortical preparatory activity. J Neurophysiol 97:348–359.

Churchland MM, Santhanam G, Shenoy KV (2006a) Preparatory activity in premotor and motor cortex reflects the speed of the upcoming reach. J Neurophysiol 96:3130–3146.

Churchland MM, Yu BM, Ryu SI, Santhanam G, Shenoy KV (2006b) Neural variability in premotor cortex provides a signature of motor preparation. Journal of Neuroscience 26:3697–3712.

Churchland MM, Cunningham JP, Kaufman MT, Ryu SI, Shenoy KV (2010) Cortical preparatory activity: representation of movement or first cog in a dynamical machine? Neuron 68:387–400.

Churchland MM, Cunningham JP, Kaufman MT, Foster JD, Nuyujukian P, Ryu SI, Shenoy KV (2012) Neural population dynamics during reaching. Nature 487:51–+.

Crammond DJ, Kalaska JF (2000) Prior information in motor and premotor cortex: Activity during the delay period and effect on pre-movement activity. J Neurophysiol 84:986–1005.

Duan CA, Pan Y, Ma G, Zhou T, Zhang S, Xu NL (2021) A cortico-collicular pathway for motor planning in a memory-dependent perceptual decision task. Nat Commun 12:2727.

Economo MN, Viswanathan S, Tasic B, Bas E, Winnubst J, Menon V, Graybuck LT, Nguyen TN, Smith KA, Yao ZZ, Wang LH, Gerfen CR, Chandrashekar J, Zeng HK, Looger LL, Svoboda K (2018) Distinct descending motor cortex pathways and their roles in movement. Nature 563:79–+.

Elsayed GF, Lara AH, Kaufman MT, Churchland MM, Cunningham JP (2016) Reorganization between preparatory and movement population responses in motor cortex. Nat Commun 7.

Finkelstein A, Fontolan L, Economo MN, Li N, Romani S, Svoboda K (2021) Attractor dynamics gate cortical information flow during decision-making (Apr, 10.1038/s41593-021-00840-6, 2021). Nat Neurosci 24:897–897.

Heitz RP (2014) The speed-accuracy tradeoff: history, physiology, methodology, and behavior. Front Neurosci 8:150.

Inagaki HK, Inagaki M, Romani S, Svoboda K (2018) Low-Dimensional and Monotonic Preparatory Activity in Mouse Anterior Lateral Motor Cortex. Journal of Neuroscience 38:4163–4185.

Inagaki HK, Fontolan L, Romani S, Svoboda K (2019) Discrete attractor dynamics underlies persistent activity in the frontal cortex. Nature 566:212–+.

Inagaki HK, Chen SS, Daie K, Finkelstein A, Fontolan L, Romani S, Svoboda K (2022a) Neural Algorithms and Circuits for Motor Planning. Annual Review of Neuroscience 45:249–271.

Inagaki HK, Chen SS, Ridder MC, Sah P, Li N, Yang ZD, Hasanbegovic H, Gao ZY, Gerfen CR, Svoboda K (2022b) A midbrain-thalamus-cortex circuit reorganizes cortical dynamics to initiate movement. Cell 185:1065–+.

Janssen P, Shadlen MN (2005) A representation of the hazard rate of elapsed time in macaque area LIP. Nat Neurosci 8:234–241.

Jaramillo S, Zador AM (2011) The auditory cortex mediates the perceptual effects of acoustic temporal expectation. Nat Neurosci 14:246–251.

Jiang X, Saggar H, Ryu SI, Shenoy KV, Kao JC (2020) Structure in Neural Activity during Observed and Executed Movements Is Shared at the Neural Population Level, Not in Single Neurons. Cell Rep 32:108148.

Kao TC, Hennequin G (2018) Null Ain’t Dull: New Perspectives on Motor Cortex. Trends Cogn Sci 22:1069–1071.

Kao TC, Sadabadi MS, Hennequin G (2021) Optimal anticipatory control as a theory of motor preparation: A thalamo-cortical circuit model. Neuron 109:1567–+.

Kaufman MT, Churchland MM, Ryu SI, Shenoy KV (2014) Cortical activity in the null space: permitting preparation without movement. Nat Neurosci 17:440–448.

Kilavik BE, Confais J, Riehle A (2014) Signs of timing in motor cortex during movement preparation and cue anticipation. Adv Exp Med Biol 829:121–142.

Kobak D, Brendel W, Constantinidis C, Feierstein CE, Kepecs A, Mainen ZF, Qi XL, Romo R, Uchida N, Machens CK (2016) Demixed principal component analysis of neural population data. Elife 5.

Kunimatsu J, Suzuki TW, Ohmae S, Tanaka M (2018) Different contributions of preparatory activity in the basal ganglia and cerebellum for self-timing. Elife 7.

Lara AH, Elsayed GF, Zimnik AJ, Cunningham JP, Churchland MM (2018) Conservation of preparatory neural events in monkey motor cortex regardless of how movement is initiated. Elife 7.

Li N, Daie K, Svoboda K, Druckmann S (2016a) Robust neuronal dynamics in premotor cortex during motor planning. Nature 532:459–464.

Li N, Daie K, Svoboda K, Druckmann S (2016b) Robust neuronal dynamics in premotor cortex during motor planning. Nature 532:459–464.

Li N, Chen TW, Guo ZV, Gerfen CR, Svoboda K (2015) A motor cortex circuit for motor planning and movement. Nature 519:51–U88.

Michaels JA, Dann B, Scherberger H (2016) Neural Population Dynamics during Reaching Are Better Explained by a Dynamical System than Representational Tuning. Plos Comput Biol 12.

Murakami M, Vicente MI, Costa GM, Mainen ZF (2014) Neural antecedents of self-initiated actions in secondary motor cortex. Nat Neurosci 17:1574–1582.

Palmer J, Huk AC, Shadlen MN (2005) The effect of stimulus strength on the speed and accuracy of a perceptual decision. J Vis 5:376–404.

Ratcliff R, Smith PL (2004) A comparison of sequential sampling models for two-choice reaction time. Psychol Rev 111:333–367.

Ratcliff R, Cherian A, Segraves M (2003) A comparison of macaque behavior and superior colliculus neuronal activity to predictions from models of two-choice decisions. J Neurophysiol 90:1392–1407.

Remington ED, Narain D, Hosseini EA, Jazayeri M (2018) Flexible Sensorimotor Computations through Rapid Reconfiguration of Cortical Dynamics. Neuron 98:1005–+.

Riehle A, Requin J (1989) Monkey Primary Motor and Premotor Cortex - Single-Cell Activity Related to Prior Information About Direction and Extent of an Intended Movement. J Neurophysiol 61:534–549.

Riehle A, MacKay WA, Requin J (1994) Are extent and force independent movement parameters? Preparation- and movement-related neuronal activity in the monkey cortex. Exp Brain Res 99:56–74.

Rosenbaum DA (1980) Human movement initiation: specification of arm, direction, and extent. J Exp Psychol Gen 109:444–474.

Sadtler PT, Quick KM, Golub MD, Chase SM, Ryu SI, Tyler-Kabara EC, Yu BM, Batista AP (2014) Neural constraints on learning. Nature 512:423–426.

Servant M, Tillman G, Schall JD, Logan GD, Palmeri TJ (2019) Neurally constrained modeling of speed-accuracy tradeoff during visual search: gated accumulation of modulated evidence. J Neurophysiol 121:1300–1314.

Shenoy KV, Sahani M, Churchland MM (2013) Cortical Control of Arm Movements: A Dynamical Systems Perspective. Annual Review of Neuroscience, Vol 36 36:337–359.

Stavisky SD, Kao JC, Ryu SI, Shenoy KV (2017) Motor Cortical Visuomotor Feedback Activity Is Initially Isolated from Downstream Targets in Output-Null Neural Stat Space Dimensions. Neuron 95:195–+.

Tanaka M (2007) Cognitive signals in the primate motor thalamus predict saccade timing. J Neurosci 27:12109–12118.

Vyas S, Golub MD, Sussillo D, Shenoy KV (2020) Computation Through Neural Population Dynamics. Annual Review of Neuroscience, Vol 43 43:249–275.

Wang J, Narain D, Hosseini EA, Jazayeri M (2018) Flexible timing by temporal scaling of cortical responses. Nat Neurosci 21:102–+.

Xu D, Dong M, Chen Y, Delgado AM, Hughes NC, Zhang L, O’Connor DH (2022) Cortical processing of flexible and context-dependent sensorimotor sequences. Nature 603:464–469.

Yang W, Tipparaju SL, Chen G, Li N (2022) Thalamus-driven functional populations in frontal cortex support decision-making. Nat Neurosci 25:1339–1352.

Zariwala HA, Kepecs A, Uchida N, Hirokawa J, Mainen ZF (2013) The Limits of Deliberation in a Perceptual Decision Task. Neuron 78:339–351.

